# Transcriptome-wide prediction of lncRNA-RNA interactions by a thermodynamics algorithm

**DOI:** 10.1101/126946

**Authors:** Ivan Antonov, Andrey Marakhonov, Maria Zamkova, Mikhail Skoblov, Yulia Medvedeva

**Affiliations:** Institute of Bioengineering, Research Center of Biotechnology RAS, Moscow, 117312, Russia; Department of Molecular and Biological Physics Dolgoprudny, Moscow Region, 141701, Russia; Laboratory of Functional Analysis of the Genome, Moscow Institute of Physics and Technology, Dolgoprudny, Moscow Region, 141701, Russia; Federal state scientific budgetary Institution “Research Centre for Medical Genetics” Moscow, 115478, Russia; Russian N.N.Blokhin Cancer Research Center, Moscow, 115478, Russia; Vavilov Institute of General Genetics, RAS, Moscow, 119333, Russia

## Abstract

**Motivation:** The discovery of thousands of long noncoding RNAs (lncRNAs) in mammals raises a question about their functionality. It has been shown that some of them function post-transcriptionally via formation of inter-molecular duplexes. Sequence alignment tools are frequently used for transcriptome-wide prediction of RNA-RNA interactions. However, such approaches have poor prediction accuracy since they ignore RNA secondary structure and interaction energy. On the other hand, application of the thermodynamics-based algorithms to long transcripts is not computationally feasible on a large scale.

**Results:** Here we describe a new computational pipeline ASSA that combines sequence alignment and thermodynamics tools for efficient prediction of RNA-RNA interactions between long transcripts. ASSA outperforms four other tools in terms of the Area Under the Curve. ASSA predictions for the lncRNA *HOTAIR* confirm that it binds to the chromatin through hybridization with the nascent transcripts. Analysis of the 49 murine lncRNA knockdown experiments reveals one transcript that may regulate its targets via RNA-RNA interactions.

**Availability:** ASSA is available at http://assa.sourceforge.net/.

## 1 Introduction

Due to the single strand nature of an RNA molecule its nucleotides are capable of base pairing with the complementary nucleotides. Usually the hybridization occurs between different regions of the same transcript producing the secondary structure. However, a part of one RNA molecule can bind to a complementary part of another RNA molecule forming inter-molecular duplex. Such RNA-RNA pairing is called *antisense interaction* and the corresponding RNAs are known as natural antisense transcripts or NATs [1].

Long noncoding RNAs (lncRNAs) are a large and diverse class of transcribed RNA molecules with a length of more than 200 nucleotides that do not encode proteins. The discovery of thousands of lncRNAs expressed in the mammalian cells raises a question about their functionality [2, 3]. The fact that the whole set of lncRNAs transcription is regulated [4], indirectly supports their functionality. Due to the functional diversity [5] the role and/or the molecular mechanism of only a few hundred lncRNAs have been determined to date. Particularly, it has been shown that some of them function post-transcriptionally via formation of inter-molecular RNA-RNA duplexes [6, 7, 8].

The primary aim of this work is to bioinformatically address novel lncRNA functions by predicting RNA-RNA interactions transcriptome-wide. Many large-scale computational studies of mammalian NATs [9, 10, 11, 12] have utilized sequence alignment tools (such as BLASTn [13] or LASTAL [14]) without taking into account the RNA secondary structure and the interaction energy, crucial for the RNA binding. To compensate for this disadvantage one can use a thermodynamics-based method to perform a co-folding of two RNAs (the query RNA versus each RNA in the transcriptome) into the minimal free energy (MFE) structure.

The major disadvantage of a thermodynamics-based approach is the computational complexity that does not allow its application to a genome- or transcriptome-wide analysis. Here we present a new pipeline, called ASSA (“AntiSense Search Approach”), which reduces running time of a thermodynamics-based search by fast identification of putative antisense sites using a sequence alignment tool BLASTn [13] followed by verification of each potential interaction by a cofolding tool *bifold* [15]. In this pipeline we (i) automate selection of the initial set of the putative antisense sites (i.e. optimize the BLASTn search thresholds), (ii) estimate the statistical significance (E-value) of the antisense interaction energy and (iii) optimize the length of the flanking sequences to the putative sites for the *bifold* run.

A similar idea has been used in a recent study by [16] where the local alignment tool LASTAL [14] has been applied to identify the initial set of “seed” sites followed by the calculation of the interaction energy by the *IntaRNA* tool [17]. However, no statistical significance has been assigned to the obtained interaction energies. This is an important issue since the “background” energies (the values produced by the random sequences) largely depend on the lengths of the input transcripts (see our analysis below).

We evaluate the ASSA performance on a set of experimentally validated functional NATs. Moreover, the comparison of our pipeline with several other large-scale computational tools demonstrates that ASSA performance is more accurate and less time consuming. Our analysis indicates that ASSA can potentially be used to search for the lncRNAs functioning via *short-trans* antisense duplexes.

## 2 Materials and Methods

### 2.1 The ASSA algorithm

Several thermodynamics-based algorithms [18, 19, 20, 21], including *bifold* [15], can be used to predict an antisense interaction with respect to the RNA secondary structure. Unfortunately it is not computationally feasible to apply these tools to the long RNAs and complete transcriptomes due to the large execution time (see Supplementary Figure 1). Thus, we develop an approximation approach, ASSA (“AntiSense Search Approach”), that identifies antisense partners for a transcript of any length on a large scale using a co-folding algorithm. To achieve this aim we perform several trainings to optimize the ASSA parameters.

As the first step of the ASSA pipeline we use the local sequence alignment tool BLASTn (with the “`-strand minus`” option) to identify the possible intermolecular duplexes. In this work, we refer to the produced local alignments as the *“putative antisense sites*”. We perform the BLASTn search with the only threshold for the seed length (required by the BLASTn) but no threshold for the E-value. By default, ASSA uses seed of the length 10 identifying all the local alignments that contain at least one perfect antisense duplex of length ≥ 10 bp. Next, to select the putative sites that are likely to be reconstructed by the *bifold* all the BLASTn hits were filtered by the alignment length, GC content and percent complementarity.

Thus, the Training 1 is performed in order to optimize the thresholds for the local alignment filtering (see Supplementary Text). For this purpose, we simulate the input to the *bifold* in an ASSA run by generating special sequence pairs. The middle part of each sequence in a pair represents a putative antisense site (a BLASTn local alignment) with a particular length, GC content and percent complementarity. The antisense site is flanked by the random sequences of length 50 nt on both sides (see Supplementary Figure 2). In total, we produce 115,500 sequence pairs corresponding to “pseudo”-BLASTn hits with various length (from 10 bp to 50 bp), GC content (from 0% to 100%) and percent complementarity (from 80% to 100%). We apply *bifold* to all the sequence pairs and calculate the fraction of the original site length that is predicted by the cofolding tool. For each specific set of GC content and percent complementarity we selecte the site length that is reconstructed by the *bifold* in at least 75% of cases. We use the linear regression to estimate the site length threshold (*SLT*) depending on its GC content and percent complementarity (see Supplementary Text).

In every ASSA run the putative antisense sites are selected according to the formula derived in the Training 1. The use of putative antisense sites significantly reduces the *bifold* execution time by applying the tool to the specific sequence chunks rather than the full-length transcripts. Namely, to compute the interaction energy of a putative site (ΔΔ*G*) *bifold* is applied to the regions of two transcripts consisting of the BLASTn hit together with the flanking sequences of specific length on both sides.

We perform the Training 2 in order to study the dependance between the lengths of the sequences submitted to the *bifold* and the produced interaction energy (see Supplementary Text). In order to do that we apply *bifold* to the 1700 pairs of random sequences with various lengths (ranging from 200 nt to 1400 nt) and compute the distributions of the obtained interaction energies (see Supplementary Figure 3). This analysis demonstrate that longer random sequences produce stronger interaction energies. For example, the average interaction energy obtained from two random sequences of length 200 nt each is -9.8 kcal/mole, while the average ΔΔ*G* value for sequences of length 1000 nt is -15.7 kcal/mole. Thus, in the ASSA pipeline the statistical significance of a ΔΔ*G* energy (P-value) is calculated with respect to the lengths of both sequences submitted to the *bifold* (i.e. the lengths of the sequence chunks involved in the local alignment plus the flank lengths). Next, for each transcript pair ASSA computes the *Interaction Score* by combining the P-values from all the putative sites found between the two RNAs. The statistical significance of the observed score is estimated using E-value (see Training 2 in the Supplementary Text).

For optimization purposes, the calculation of the ΔΔ*G* values in the ASSA pipeline is performed by gradually increasing the length of the flanking sequences and running *bifold* on each iteration. We perform the Training 3 to optimize the lengths of the flanks used in this iterative process (see Supplementary text). We use the *TINCR* pull-down experiment [22] to prepare a training set which consists of the top 200 transcripts identified in the experiment and the 600 randomly chosen human RNAs. ASSA is applied to the training set with different parameters for the *bifold* loop. After each iteration the prediction accuracy is measured by the Area Under the Curve (AUC) value. Unexpectedly, usage of the flanks longer than 50 nt frequently decreases the prediction accuracy (see Supplementary Figure 4). Based on this analysis the default values for the `‐‐flank_min`, `‐‐flank_step` and `‐‐flank_max` ASSA options are assigned to 25 nt, 25 nt and 50 nt, respectively. It should be noted that ASSA merges any two adjacent putative sites that overlap by their flanks on both transcripts. If putative sites overlap on one sequence only, they are considered to be separate sites.

### 2.2 All-vs-all BLASTn search and repeat masking

Information about the genes expressed in the K562 cell line was taken from the FANTOM5 database (sample id “CNhs12334.10824-111C5”). All the genes with non-zero expression value were considered. For the alternatively spliced genes the longest transcript was selected only.

The “all-vs-all” BLASTn search was performed in the antisense mode with the seed length equal to 15 and without a threshold on the E-value. Thus, an antisense interaction was recorded between ant two transcripts with at least one perfectly complementary duplex of length 15 bp or more. This relatively strict value for the BLASTn seed threshold was chosen taking into account that even AT-rich duplexes of this length were frequently reconstructed by the *bifold* (see Supplementary Fig.2). The number of the antisense partners for a given RNA was defined as the number of the unique transcripts. Note, that several BLASTn local alignments could be found between two sequences. Thus, the total number of the antisense partners was less than the total number the BLASTn hits.

We ran the RepeatMasker in a quick mode (the `-qq` option) to mask the human-specific repeats (`-species human`) in all the sequences. Additionally, the `-alu` option was used to restrict masking to the *Alu* repeats only.

### 2.3 Prediction of the RNA-RNA and RNA-DNA interactions

To predict the RNA-RNA interactions we ran ASSA, RRP and LncTar with the default parameters; LASTAL – with the custom substitution matrix and the parameters from [12]: `-a20 -b8 -s0 -e1`; the IntaRNA – with the parameters recommended by the authors [23]: `-p 7 -w 150 -L 100`. The RRP predictions were sorted according to the “SUMENERGY” values.

Triplexator was applied to search the whole human genome (hg19) with the settings recommended at the official web-site: `-l 15 -e 20 -c 2 -fr off -g 20 -fm 0 -of 0 -rm 2`. The Triplexator score for each genomic region was calculated as the sum of the scores of all the predictions inside the region.

### 2.4 Analysis of the lncRNA knockdown in murine cells

Since we are focusing on the RNA-RNA interactions it is reasonable to search for the lncRNA antisense interactions only within the genes expressed in the mESCs. To compile such a list we take advantage of the large number of the various knockdown experiments performed by [24] – there are 187 experiments in total (147 lncRNA and 40 transcription factor knockdowns). We hypothesize that a gene, which has never been down-regulated in a wide range of the knockdown experiments, is probably not expressed in the corresponding cell line. Thus, we select the genes that have been significantly down-regulated in at least one (out of the 187) knockdown experiment performed in the study. There are 3695 such genes represented by the 5451 transcripts.

Among the 147 lncRNAs provided by [24] there are 100 sequences containing “N” characters which can not be used as ASSA input. Thus, we used the BLASTn (with the seed length = 40) to match the 147 transcripts with the recent GENCODE annotation (release 19). We managed to resolve the 29 “N”-containing lncRNAs. The remaining 71 noncoding transcripts with ambiguous nucleotide(s) were not used for further analysis. Additionally, we discarded 19 lncRNAs with less than 50 differentially expressed genes (among the 3695 genes) as well as 8 *Alu*-containing sequences. This resulted in a final list of 49 lncRNAs.

ASSA was run without a threshold on E-value since we wanted to sort the 3695 genes by their ability to hybridize with the query sequence rather than identify the top predictions only. The GSEA FDR value was computed using the *gsea2.jar* tool with 10000 permutations. To obtain the random sequences, we used the *uShuffle* tool [25] for di-nucleotide shuffling, while the mono-nucleotide shuffling was performed using a custom Perl script.

## 3 Results

### 3.1 The ASSA pipeline

Here we present a new computational pipeline, called ASSA (“AntiSense Search Approach”), developed for the thermodynamics-based prediction of the RNA-RNA interactions on a transcriptome scale. The general idea of this approach is to first identify putative antisense sites by the local sequence alignment tool BLASTn [13], compute the interaction energy (ΔΔ*G*) for each site by the RNA co-folding tool *bifold* [15] and using the obtained ΔΔ*G* values estimate the statistical significance of each predicted RNA-RNA interaction (see Fig.1).

**Figure 1:**
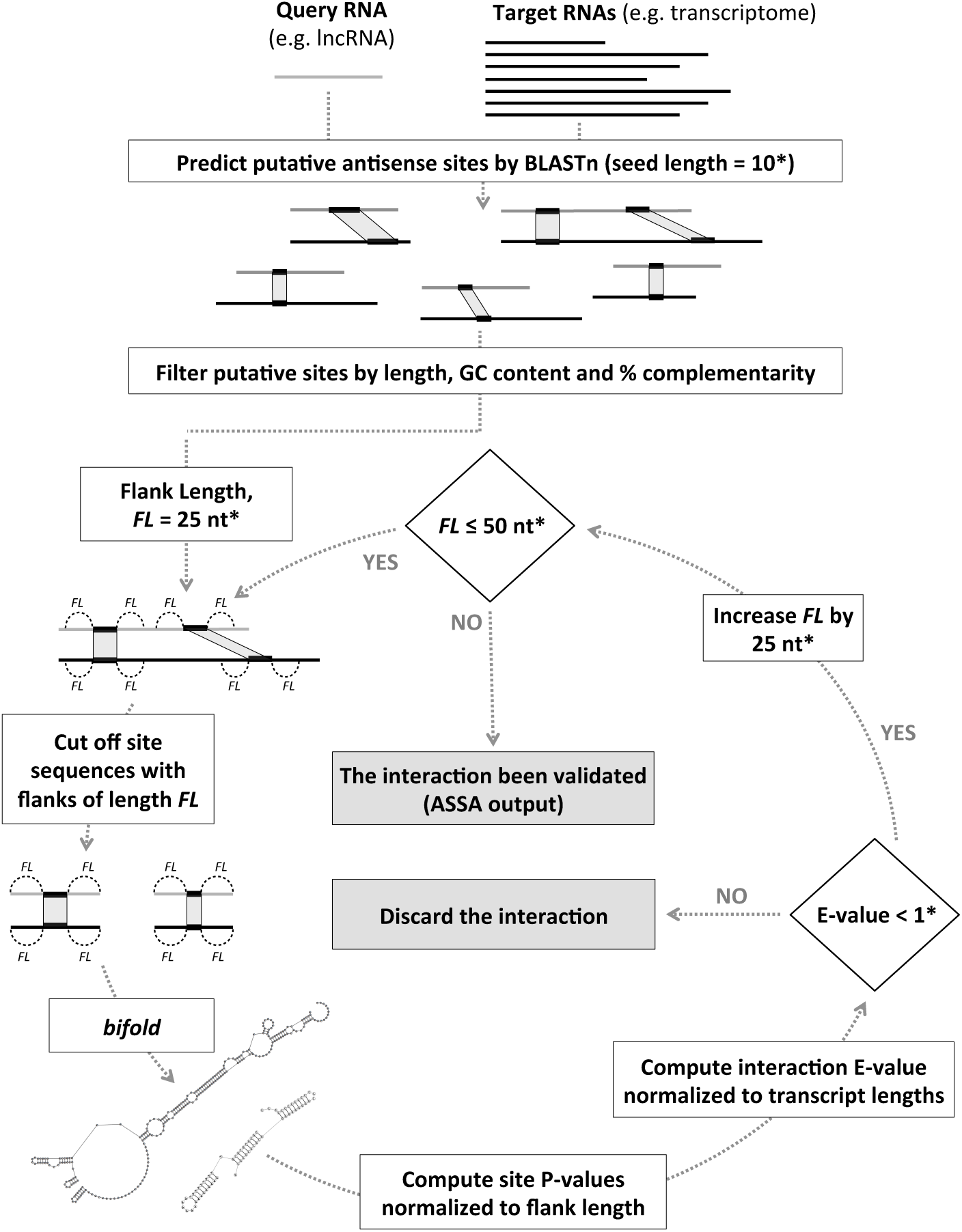
The ASSA pipeline. *The parameters can be adjusted by the ASSA options (the default values are shown on the figure).

ASSA takes two sets of nucleotide sequences as input (the query and the target transcripts). On the first step, the antisense mode of the BLASTn is used to search all the query RNAs versus all the targets. The obtained local alignments are filtered by the length, GC content and percent complementarity producing a set of putative antisense sites (see Material and Methods).

On the next step, ASSA uses the *bifold* tool to compute the interaction energies (ΔΔ*G*) for all the sites. It should be noted that the ΔΔ*G* values are computed from the transcript regions containing the putative sites together with the flanking sequences on both sides. The obtained energies are used to estimate the statistical significance (measured by E-value) of the interaction between any two transcripts that have at least one putative antisense site. Transcript pairs that do not have antisense sites or with E-values that do not satisfy the user-defined threshold are discarded.

The length of the flanks that are added to the putative antisense sites is an important parameter of the ASSA pipeline. On the one hand, longer flanks allow to take into account more elements of the RNA secondary structure. On the other hand, the *bifold* execution time increases with the length of the flanking sequences. Thus, in the ASSA pipeline we iteratively increase the value of the flank length parameter and perform the ΔΔ*G* calculation and the hits filtering on every iteration (see Fig.1). The default values of the iteration process parameters were chosen based on the ASSA application to a training set (see Materials and Methods). By default, the flank length is increased from 25 nt to 50 nt in two iterations.

ASSA outputs a list of predicted RNA-RNA interactions sorted by E-value. It should be noted that the HTML-based output which conveniently visualizes the location of the predicted antisense duplexes can also be produced (installation of additional Perl modules is required to enable this option).

ASSA execution time depends on the query transcript length, the size of the target database and the length of the flanking sequences. The RNA co-folding is the most time consuming step of the pipeline. Since *bifold* is independently applied to each putative site, this process could easily be distributed between several CPUs (using the `‐‐num_threads` option) to speed up the ASSA run.

### 3.2 Classification of the antisense interactions

Our literature search for the published cases of the biologically active duplexes formed between long RNAs (i.e. mRNA-mRNA, lncRNA-mRNA or lncRNA-lncRNA) in mammals prompted us to expand the classical “cis-trans” classification of the natural antisense transcripts.

Antisense interactions are usually classified into two groups – *cis* and *trans*. The interactions of the first type occur between the products of the overlapping genes that are transcribed in opposite directions. The resulting RNAs have one or several (due to splicing) sites that are perfectly complementary to each other. All other interactions are classified as *trans* (“not-cis ”) since they are formed between the transcripts of genes located in different genomic regions.

A special type of the *trans* RNA-RNA binding occurs when one of the overlapping genes has an expressed copy (ortholog) at another locus. The duplicate of the gene harbors a sequence highly complementary to a part of the other gene in the overlap and thus can form *trans*-antisense duplexes with it. This scenario has been observed in the case of expressed pseudogenes [26]. Moreover, it has been shown that such pseudogene related duplexes can be recognized by RNAi machinery and produce functional siRNAs in mouse oocytes [27]. Inversion of a genomic region during gene duplication is another possible scenario for such NATs formation [28]. So, we refer to these interactions as “*pseudo-cis*” because the transcript regions that form the inter-molecular duplex share a common ancestral sequence.

Sequence repeats of several types occupy a significant portion of the mam-malian genomes. It is a common practice in the large scale computational studies to mask repeats of all types prior to the antisense search [11, 9, 10] in order to avoid the prediction of the large number of interactions based on sequence repeats. To study the contribution of different repeat types to the total number of possible antisense partners of a given transcript we performed “all-vs-all” BLASTn search for the 10664 genes expressed in the K562 cell line. As expected the number of the predicted antisense partners linearly depended on the query length. Surprisingly, we observed three types of RNAs on the Fig. 2A. We were curious whether the origin of this observation could be related to a specific repeat family.

**Figure 2:**
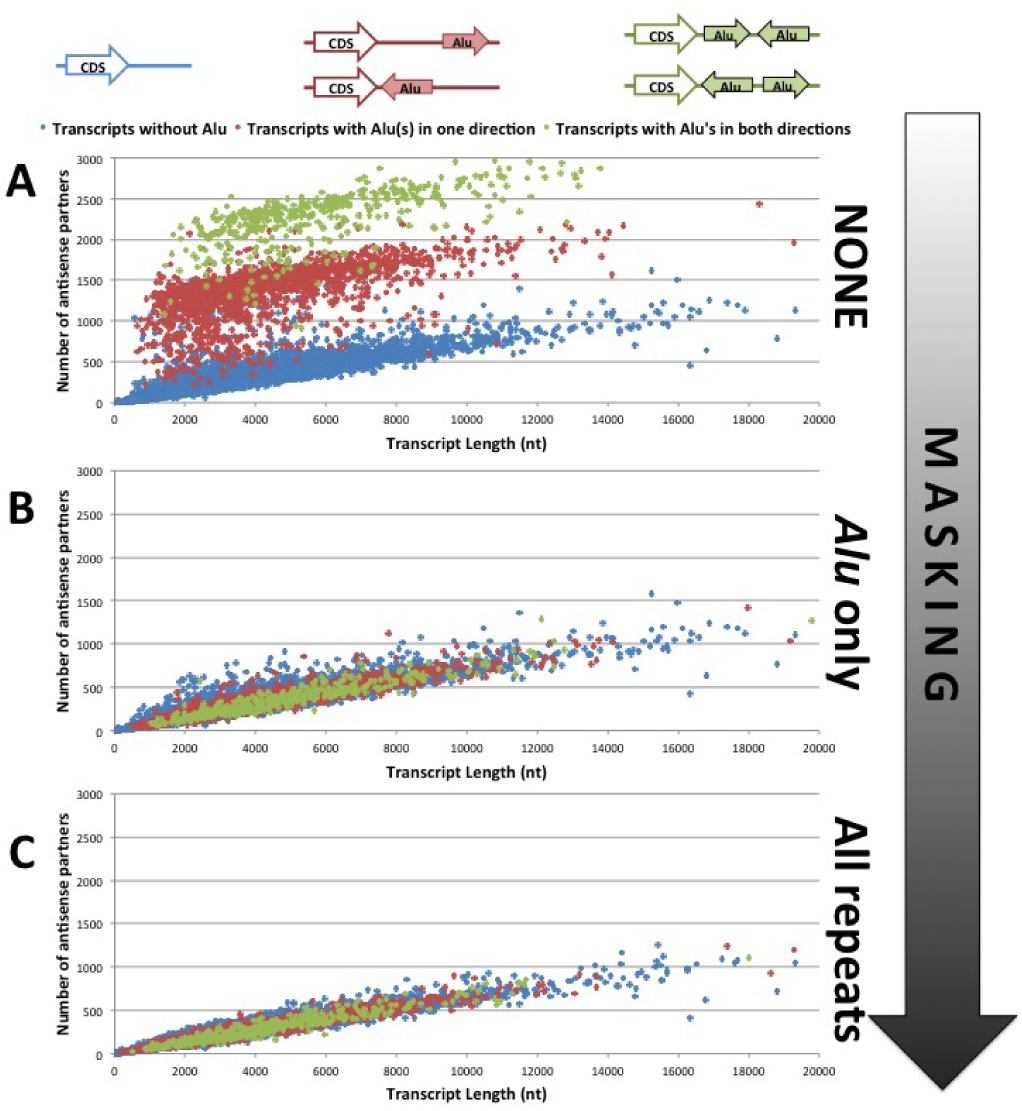
*Alu* repeats are the major contributors of the repeat-based antisense interactions between human transcripts. Dependence between the query transcript length and the number of antisense partners is shown for three types of masking: (A) no masking, (B) masking of the *Alu*-repeats only and (C) masking of all repeats (as identified by the RepeatMasker). Each dot on the graph corresponds to one transcript. The transcripts without *Alu* repeats are blue; the transcripts with all *Alu*(s) in one orientation are red; the transcripts with at least one *Alu* repeat in the direct orientation and at least one *Alu* repeat in the reverse orientation are green.

The *Alu*-repeats are up to 350 nt long and have relatively high percent identity (> 70%). They are localized in the human transcripts either in the direct or in the reverse-complement orientation. It has been shown that a pair of RNAs with *Alu* repeats in opposite directions are able to interact with each other and trigger Staufen Mediated Decay [29, 30] and/or regulate mRNA translation [31]. We hypothesized that the three groups of transcripts observed on Fig.2A were related to the *Alu*-contaning transcripts.

To check this hypothesis we applied the RepeatMasker [32] to the 10664 transcripts and identified 2212 *Alu*-containing sequences. Thus, on Fig.2 we used different colors for (i) the 8452 transcripts without *Alu* repeats, (ii) the 1807 RNAs with *Alu*(s) in one direction only and (iii) the 405 RNAs with at least one *Alu* repeat in the direct and at least one repeat in the opposite orientation. This color-scheme matched well with the three groups (see Fig.2A). Indeed, the transcripts with two or more *Alu*-repeats in different directions are able to form inter-molecular duplexes with any other *Alu*-containing transcript. RNAs with *Alu*(s) in one orientation can only hybridize with transcripts containing complementary repeat sequence. Clearly, transcripts without *Alu*’s have the lowest antisense potential since they can not participate in the *Alu*-based interactions.

To further confirm the observed role of the **Alu** repeats we masked them in the 2212 sequences and repeated the “all-vs-all” search for all the 10664 transcripts. The same linear dependence was observed for all the queries (Fig. 2B) proving that the presence of an *Alu* repeat(s) in a query significantly increased its antisense potential.

Finally, we used the RepeatMasker once again to mask repeats of all types (7.8% of the total sequence length) and repeated the “all-vs-all” search. We did not observe a significant change of the graph (Fig. 2C). This results suggests that the *Alu* repeats play the major role in the repeat-based interactions in the human cells. We, thus, define yet another type of *trans*-antisense interactions – the “*Alu*-based”.

We call *short-trans* any interaction that is not *cis* and does not have *pseudo-cis* or *Alu*-based duplexes. Thus, we define four types of NATs – *cis, pseudo-cis, Alu*-based and *short-trans*.

### 3.3 ASSA predictions for the functional NATs

In order to evaluate ASSA performance we applied it to the 34 functional natural antisense transcripts (NATs) collected from the literature – 11 *cis*, 4 *pseudo-cis*, 10 *Alu*-based and 9 *short-trans* cases (see Supplementary Table 1). ASSA predictions for *cis, pseudo-cis* and *Alu*-based interactions were very strong due to the existence of long (>100 bp) duplexes with high percent of complementarity.

By contrast, only 6 out of the 9 *short-trans* interactions were identified by ASSA and only 3 out of 6 had E-value < 0.01. The inability of ASSA to identify some of the *short-trans* cases could be due to our approach to apply *bifold* to the regions containing putative antisense sites instead of analyzing the full length transcripts. To test this possibility we applied *bifold* directly to the full-length sequences from the four *short-trans* antisense pairs with weak ASSA E-values (lncRNA-ATB::IL11, GAS5::MYC, lincRNA-p21::JUNB and lincRNA-p21::CTNNB1). Additionally, each lncRNA was used to generate 20 random sequences via mono-nucleotide shuffling and the *bifold* was applied to predict the interactions between the random lncRNAs and the corresponding mRNAs. For each of the four *short-trans* interactions the predicted duplex lengths and the ΔΔ*G* values obtained for the random sequences were similar to what was observed for the actual lncRNAs (Supplementary Figure 5). This result indicates that the inability of ASSA to detect the interactions between these *short-trans* pairs was not due to the restriction to run *bifold* on the putative site containing regions only. Thus, all four functional *short-trans* NATs did not produce statistically significant energies even if the full-length transcripts were used. This may indicate that the spontaneous hybridization of the corresponding RNAs is unlikely and the mediator proteins may be required to facilitate their binding.

According to the obtained results ASSA is suitable to predict all types of antisense interactions between long RNAs. However, the identification of the *short-trans* interactions is the most challenging task.

### 3.4 Comparison of ASSA with other tools

We compared ASSA performance with the four RNA-RNA prediction tools – IntaRNA [17], RRP [16], LASTAL [14, 12] and LncTar [33]. To evaluate the ability of the tools to predict the *short-trans* interactions we used the data from the “RIA-Seq” experiment (RNA interactome analysis, followed by deep sequencing) performed for the lncRNA *TINCR* [22]. The authors have demonstrated that *TINCR* is bound to the transcripts of 1794 genes in the cytoplasm of human keratinocytes. Additionally it has been shown that at least some of the identified RNAs are associated with *TINCR* through direct *short-trans* antisense duplexes.

From the 1794 genes we removed transcripts with *Alu* repeats (since we were focusing on the *short-trans* interactions) as well as 200 RNAs from the training set that was used for flank length optimization (see Materials and Methods). The remaining 1199 *TINCR* pull-down transcripts were used to prepare a test set containing 4796 sequences – the 1199 pull-down RNAs and 3597 (3 * 1199) randomly selected human transcripts.

Each of the five tools was used to rank the sequences from the test set according to their ability to hybridize with the 3.7 kb *TINCR* transcript (NR_027064). It should be noted that the IntaRNA required more than 8 Gb of RAM to predict *TINCR* interactions with transcripts longer than 3 kb. Due to the hardware restrictions we were able to apply IntaRNA to the subset of sequences not exceeding 3 kb in length (2846 transcripts). ASSA outperformed all the tools in terms of the Area Under the Curve (AUC) on the full test set as well as on the “≤ 3 kb” subset (see Table 1 and Fig. 3A).

**Table 1:** Performance of different tools on the full TINCR test set (4796 transcripts in total) and the subset consisting of the sequences with the length ≤ 3kb (2846 transcripts).

| Tool | Time (min) | # predictions | AUC (full) | AUC ($\leq 3$kb) |
| --- | --- | --- | --- | --- |
| ASSA | 430 (94*) | 4431 | **0.646** | **0.620** |
| IntaRNA | 23901** | 2846** | - | 0.612 |
| RRP | 1135 | 4796 | 0.636 | 0.587 |
| LASTAL | 0.07 | 3979 | 0.605 | 0.553 |
| LncTar | 2535 | 4708 | 0.464 | 0.534 |

**Figure 3:**
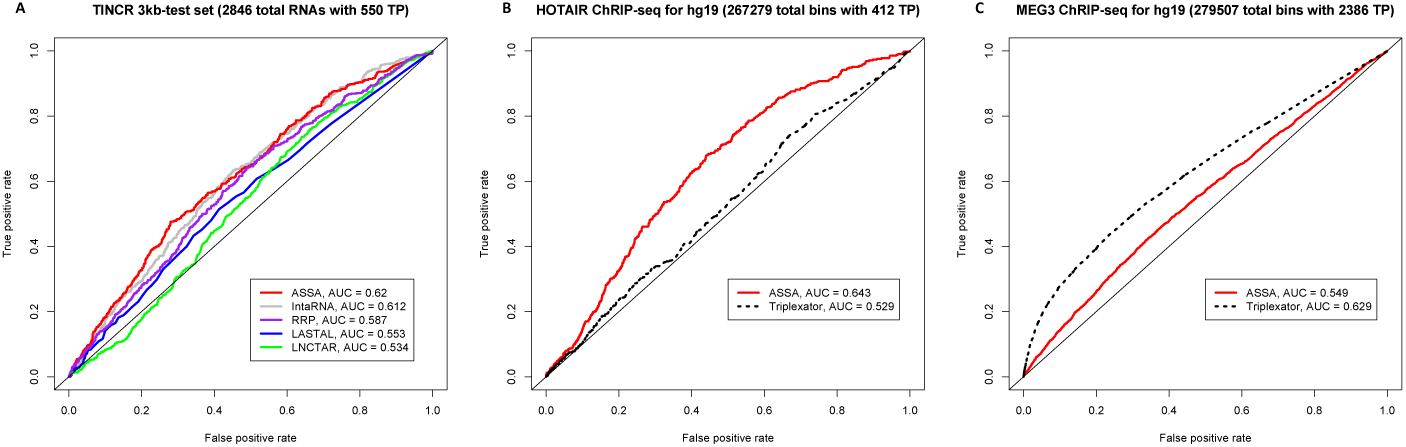
(A) The ROC curves produced by five tools for the TINCR 3kb-test set. (B,C) ASSA results support the RNA-RNA based mechanism of the lncRNA *HOTAIR* chromatin targeting. ASSA outperforms Triplexator in the accuracy of genome-wide ChIRP-seq peaks prediction for the lncRNA *HOTAIR* (B), but not for the *MEG3* (C).

It should be noted that ASSA was faster than the other thermodynamics-based tools (IntaRNA, RRP and LncTar). The sequence alignment algorithm LASTAL had the lowest execution time. Interestingly, the nucleotide substitution parameters optimized for predicting RNA-RNA interactions [12] allowed LASTAL to outperform LncTar.

To estimate the dependence between the predicted interaction strength (measured by the −*log*10(ASSA E-value)) and the transcript enrichment observed in the pull-down experiment (as reported by [22]) the Pearson correlation coefficient was computed. We observed small, but statistically significant positive correlation (*r* = 0.142; *p* < 10^−5^ – see Supplementary Figure 6).

Moreover, we analyzed another pull-down experiment performed for the *lncRNA-ATB* [34]. The four fastest tools failed to produce reasonable ROC-curves (the best AUC value of 0.538 was produced by the LncTar – see Supplementary Figure 7), suggesting that the RNA-RNA binding observed in the experiment could be non-specific or indirect.

Our analysis demonstrates that ASSA outperforms other tools in predicting short-trans RNA-RNA interactions.

### 3.5 ASSA was able to discriminate different mechanisms of RNA-chromatin interactions

It has been reported that many human long noncoding RNAs are localized in the nucleus where they can bind to various protein complexes and/or participate in chromatin remodeling [35]. The lncRNA *HOTAIR* is one of the best studied transcripts of this type [36]. It has been demonstrated that the *HOTAIR* plays a crucial role in promoting breast cancer metastasis by binding to the Polycomb repressive complex 2 (PRC2) and directing it to specific genomic loci [37]. Even though, the *HOTAIR* genome-wide target sites have been known for some time [38], the underlying binding mechanism remained unclear until the recent study by [39]. The authors have demonstrated that the *HOTAIR* forms *short-trans* RNA-RNA interactions with nascent transcripts and that the RNA matchmaker protein A2/B1 is required for this process. The direct hybridization between the *HOTAIR* and a region of the *JAM2* RNA has been validated *in vitro*.

First, we used ASSA to predict the interaction between the *HOTAIR* and the full-length pre-mRNA of the *JAM2* gene. Even though the exact duplex reported by Meredith *et al*. was not predicted (probably, due to the seed length threshold in the BLASTn step of ASSA), another region of inter-molecular binding was found in the 5’ end of the *JAM2* transcript. Thus, a statistically significant interaction (E-value = 0.00088) was predicted.

Next, we used ASSA to identify the RNA-RNA interactions of the lncRNA *HOTAIR* on the genome scale. For this purpose we analyzed the 832 sites of *HOTAIR*-chromatin interactions (peaks) identified in the ChIRP-seq experiment [38]. To run ASSA, the whole human genome was split into non-overlapping regions (“*bins*”) of length 3 kb. Bins corresponding to the poorly mappable genomic regions or areas with ambiguous “N” characters were discarded. This resulted in the 322,959 genomic bins (429 of them contained *HOTAIR* interaction sites). ASSA was used to search the *HOTAIR* transcript against the 322,959 bins producing predictions for 267,279 bins (412 of them contained *HOTAIR* interaction sites). It should be noted that ASSA searched both strands of every bin and the best E-value was selected. The obtained results were visualized as a ROC-curve (Fig.3B). The obtained AUC value (0.643) was similar to what was observed for the *TINCR* RIA-seq dataset supporting the RNA-RNA interaction mechanism for the *HOTAIR* on the genome-wide scale.

Formation of the RNA-DNA triplexes is another common mechanism that is utilized by lncRNAs to direct chromatin modifying complexes to specific genomic locations [40]. Up until now, the *HOTAIR* was not reported to form triplexes with DNA. To check whether the ASSA predictions were mechanism-specific, we used the Triplexator tool [41] to predict the RNA-DNA interactions between the *HOTAIR* and the 267,279 genomic regions according to the Hoogsteen and reverse Hoogsteen base pairing rules. The AUC value (0.529) calculated based on the Triplexator output was very close to the value of random model (0.5) suggesting the absence of triplexes between the *HOTAIR* and its target genomic regions.

As a negative control for the ASSA predictions we analyzed the ChIRP-seq peaks of the lncRNA *MEG3* which is known to hybridize with the DNA via triplex structures [42]. For the *MEG3* transcript ASSA produced predictions for 279,507 bins. As expected, Triplexator outperformed ASSA in terms of the AUC value (0.629 vs 0.549) when it was applied to the same set of genomic regions (Fig.3C).

Thus, we demonstrated that analysis of ChIRP-seq data by ASSA and Triplexator could be used to reveal the underlying molecular mechanism of RNA-chromatin interactions.

### 3.6 Predicting regulatory ***trans***-NATs from knockdown experiments

Gene knockdown experiments followed by transcriptome-wide identification of differentially expressed genes (either by microarray or RNA-seq) are frequently used to study various aspects of gene regulation. Such analyses have been applied to a number of human and murine lncRNAs. In each experiment genes that significantly changed their expression upon lncRNA knockdown have been identified. It is reasonable to assume that at least some of the differentially expressed genes are directly regulated by the corresponding lncRNA. Particularly, [24] has performed knockdown of the 147 lncRNA in the mouse embryonic stem cells (mESCs). In each experiment the genes that changed their expression by at least 2-fold have been identified using microarrays. However, for the majority of the noncoding transcripts the molecular mechanism of the observed regulation has remained unclear and no analysis has been performed with respect to the possible antisense interactions.

We used ASSA in order to find the lncRNAs that could regulate target genes via *short-trans* antisense interactions. For this analysis we selected 49 out of the 147 noncoding transcripts (see Material and Methods) and used ASSA to search each of them against 3695 genes expressed in mESCs. After each ASSA run all the genes were sorted by the E-value. The statistical significance of the enrichment of the differentially expressed genes at the top of the sorted gene list was estimated using GSEA [43]. As a negative control, we applied mono- and dinucleotide shuffling to the 49 lncRNAs generating two sets of random sequences and repeated the above analysis for them (Supplementary Table 3). Only one noncoding RNA demonstrated statistically significant enrichment – the GSEA FDR value for the *linc1607* (the corresponding RefSeq gene is *1700001G11Rik*) was < 0.0005 (Supplementary Figure 8) while the lowest GSEA FDR value observed among all the shuffled sequences was much weaker (0.01345). This result suggests that the *linc1607* may regulate its targets via direct *short-trans* antisense interactions.

## 4 Discussion

In this work we present a new computational pipeline ASSA that uses the thermodynamics algorithm *bifold* for a transcriptome-wide prediction of RNA-RNA interactions.

For the accurate prediction of an RNA-RNA interaction, the sequence alignment is not enough. Relative free energy of the antisense duplex (ΔΔ*G*) contributes to the predictive power of the algorithm a lot due to the existence of the RNA secondary structures. In our pipeline we use the RNA co-folding algorithm *bifold* that takes two RNA sequences and folds them simultaneously (’“cofolding”) into the minimal free energy (MFE) structure allowing inter-molecular base-pairing. The ΔΔ*G* value is obtained by comparing the energies of the initial (two separate RNA molecules with their own secondary structures) and the final (two RNAs with both inter-molecular duplexes and secondary structure regions) states of the system.

ASSA computes the ΔΔ*G* value to measure the strength of the hybridization between two RNAs and estimates its statistical significance with respect to the lengths of both transcripts using E-value. Sorting the interactions by the Evalue rather than the free energy allowed ASSA to outperform other tools in the accuracy to predict *short-trans* RNA-RNA interactions. Moreover, ASSA is > 50 times faster than the IntaRNA and > 2 times faster than the RRP. This made it possible to use ASSA for the genome-wide analysis of the ChIRP-seq data and for the transcriptome-wide analysis of the 49 murine lncRNAs.

Clearly, ASSA pipeline is still a simplification of the actual process that happens in the cell. In fact, the “co-folding” approaches, such as the *bifold*, may produce prediction errors predicting *trans*-interactions. The reason for this is that a *trans*-RNA-RNA interaction occurs between two transcripts with already established secondary structures. Thus, this is a dynamic process that requires unfolding of specific transcripts’ regions followed by formation of inter-molecular duplexes. This simulation can be performed by the methods of molecular dynamics. However, such approaches are even more time consuming than the RNA co-folding tools and at the moment we do not see a way to use them for the large scale analyses. Another source of errors in ASSA predictions may arise from the fact that only specific regions of the transcripts are analyzed. Indeed, it is known that RNAs include a number of long-range secondary structure interactions that are ignored by ASSA [44].

Nevertheless, our analysis of the *TINCR* RIA-seq and the *HOTAIR* ChIRP-seq datasets demonstrates that even this simplified approach is able to produce meaningful results. Interestingly, none of the bioinformatics tools managed to detect direct RNA-RNA interactions in another pull-down experiment performed for the *lncRNA-ATB*. Thus, even if a pull-down experiment identifies transcripts bound to an RNA of interest, it still remains unclear whether the observed binding is direct (RNA-RNA interaction) or indirect (e.g. occurs via an RNA binding protein). Our analysis indicates that this question can be clarified by applying RNA-RNA prediction tool(s) to the produced data and visualizing the obtained predictions as an ROC-curve.

In this work, we also suggest a classification for the NATs based on the origin of the hybridizing sequences. Namely, we define four types of antisense interactins – *cis, pseudo-cis, Alu*-based and *short-trans*. Importantly, we demonstrate that among all types of sequence repeats in the human genome, only *Alu* repeats have a striking influence on the ability of a transcript to base pair with other RNAs in the transcriptome [29, 30]. It should be noted, however, that two opposite direction *Alu* repeats that are located on the same RNA may interact with each other (intra-molecular hybridization) producing a special secondary structure element called “inverted repeat *Alu* elements” or IRAlus [45]. Such transcripts are unlikely to form inter-molecular *Alu*-based interactions. Since we did not take this into account in our analysis of repeat-based interactions the number of antisense partners for some IRAlus transcripts could be incorrect.

LncRNA-RNA hybridization may form yet another layer in the gene regulatory network. It should be noted that the prediction of ASSA or any other computational tool alone is not sufficient to make conclusions about the functionality of the interaction. There are many other factors that are critical for the predicted binding, including the cellular localization of the RNAs as well as, in some cases, the presence of specific RNA binding proteins. Thus, the search for new biologically active NATs is a more complex task than just prediction of antisense partners. Nevertheless, we believe that the improvement of the RNA-RNA interaction prediction methods is a necessary step in this direction.

## Acknowledgements

We would like to thank Andrey Leontovich for suggestions regarding aspects of statistical analysis; Mark Borodovsky for the help in obtaining the RRP tool; Alexandra Filatova, Victoriya Serzhanova and Nikolay Zernov for useful discussions about the the biological role of the long noncoding RNAs; Yuriy Shkandybin for the technical computational assistance.

## Funding

The work of I.A. was supported in part by the Dynasty Foundation Fellowship [No DP-B-26/14 to I.A.]; the work of Y.M was supported in part by the Russian Science Foundation grant [14-15-30002 to Y.M.].

